# Bidirectional links between food intake and sleep: a naturalistic mobile EEG study

**DOI:** 10.64898/2026.08.31.748421

**Authors:** Lilla Kovács, Péter Przemyslaw Ujma

**Affiliations:** Semmelweis University, Institute of Behavioural Sciences

**Keywords:** food intake, chrononutrition, mobile EEG, sleep

## Abstract

Sleep and nutrition are cornerstones of health, with likely reciprocal links between the two. The bidirectional link between sleep and food intake has been extensively studied, however, this literature relies almost completely on interventional and epidemiological studies. In the current study, we studied this link under naturalistic conditions, using data from a large sample of healthy participants. We investigated the bidirectional, time-lagged, within-person covariation between sleep, recorded by mobile EEGs and morning questionnaires; and food intake, recorded by retrospective evening questionnaires about the timing, content and satiation of previous-day meals. The findings supported links between sleep and food intake in both temporal directions. Later timing of the last meal was associated with later sleep onset, while a longer fasting period between the last meal and sleep was associated with reduced total sleep time, and reduced sleep efficiency. Sleep efficiency was higher following higher total daily energy intake. Later sleep was associated with a shift of next-day food intake to later hours. Overall, this naturalistic study suggests that the bidirectional relationship between sleep and food intake is primarily related to timing, not quality or composition. Contrary to some recommendations, we found no negative effect of reduced sleep quality after large, late, or satiating last meals.

## Introduction

Sleep and nutrition are key modifiable determinants of human health, both of which are independently associated with metabolic, cardiovascular, and psychiatric disorders [1–3]. Although much of the literature considers sleep and nutrition as separate determinants of health, more recent research suggests that they are tightly interconnected, with implications for understanding and modifying health-related behaviors. The relationship between nutrition and sleep is potentially bidirectional, with both food intake influencing sleep and sleep influencing food intake the following day.

In the first direction, food intake may influence sleep via either biochemical or psychological mechanisms. Biochemical mechanisms may involve the consumption of foods rich in compounds involved in the serotonergic pathway with a possible sleep-promoting effect. These include tryptophan (e.g., milk whey [4]), melatonin (e.g., tart cherries [5]), or serotonin (e.g., kiwifruit [6]). Because the metabolism of these compounds is complex and the production of sleep-promoting agents may depend on other nutrients, other aspects of the dietary profile (e.g., Mediterranean diet, glycemic index) may interact with the effects of potentially sleep-promoting foods [7]. Other possible mechanisms include the inhibition of wake-promoting hypothalamic orexin neurons by glucose, contributing to postprandial sleepiness [8], or the release of sleep-promoting cholecystokinin from the gut following fat or protein consumption [9]. In line with these proposed mechanisms, reviews of the literature have linked protein- and carbohydrate-rich diets to alterations in sleep duration or quality [10,11]. Conversely, psychological mechanisms may involve a reward or temporal cue function of food intake. Motivation and reward systems affect vigilance and may be activated by the anticipation or consumption of food, affecting subsequent sleep and wakefulness [12]. Furthermore, food intake is an important temporal cue, capable of influencing the circadian regulation of at least peripheral clocks, or of producing „masking effects”, that is an immediate change in sleep-wake patterns without modifying internal clocks [13]. A combination of reward and temporal cue mechanisms may occur if food consumption becomes a conditioned temporal cue [14]: for example, in a person accustomed to eating a large meal before sleep, the repeated association between the meal and bedtime may eventually contribute to a lack of sleepiness unless the meal is consumed.

In the other direction, sleep may influence next-day food intake through various, non-exclusive mechanisms [7]. First, shorter sleep may leave time for a longer feeding window, resulting in more opportunity for dietary intake [15,16]. Second, short or insufficient restorative sleep may interfere with appetite-regulating hormones, including leptin, ghrelin and orexin [17]. Third, poor sleep may have a psychological effect, involving an increased valuation of energy-rich food as a reward, including changes in taste perception [18] and a preference for low-quality, sugar-rich food [19,20].

Evidence for the bidirectional link between sleep and food intake has been extensively reviewed and meta-analyzed [7,10,11,21–25]. While some effects - for example, positive effects of carbohydrate- or protein-rich diets on sleep [10], or negative effects of sleep restriction on subsequent energy intake and dietary quality [23,24] - have received support, the literature is not unequivocal [10,25].

A limitation of the extant literature on sleep and food intake is that virtually all evidence comes either from interventional studies that often drastically and artificially alter sleep or nutritional patterns (e.g., [26]), or epidemiological studies that analyze correlations between dietary and sleep patterns (e.g., [27]). Despite their merits and widespread adoption, both approaches have limitations in uncovering bidirectional effects between sleep and food intake which are relevant to everyday life. In interventional studies, the complexity of the protocol often leads to small participant groups and low statistical power, and the nature of the intervention (e.g., drastic sleep restriction or extreme diets) limits ecological validity. Conversely, in epidemiological studies causal inference is limited by their correlational design.

Multiday observational studies [28] also known as naturalistic prospective studies [29] or micro-longitudinal studies [15] are an alternative approach to studying the interaction between sleep and daytime experiences. These studies implement a naturalistic design in which participants record both sleep and daily experiences over an extended period, allowing researchers to examine naturally occurring, time-lagged, within-person covariation to uncover bidirectional links. Despite the widespread use of this design to study the interaction of sleep with, for example, mood [29], daytime experiences [30], or exercise [31], we are aware of only a single study with such a naturalistic design to explore the bidirectional link between sleep and food intake. Hoopes et al. 2023 [15] studied the natural covariation of wristwatch-recorded sleep and photograph-assisted dietary records in 63 participants who provided data for 14 days. In the most informative within-participant analyses, the study primarily found a temporal cue effect in both directions, with later eating associated with later sleep, and vice versa. However, the use of actigraphy instead of EEG limited the exploration of fine-grained changes in sleep structure.

In the current study, we investigated the bidirectional, within-person covariation of sleep and food intake under naturalistic conditions. Our study aimed to bridge key gaps in the previous literature by using a large sample of 267 participants to overcome the limited statistical power of interventional studies; time-lagged, within-participant data to reduce confounding by time-invariant person-level characteristics in epidemiological research [29]; and mobile EEG and detailed daily food diaries to provide a more comprehensive assessment of sleep and food intake.

## Methods

### Data

We used data from the Budapest Sleep, Experiences and Traits Study (BSETS). The full protocol of BSETS has been published separately [28], and the description in this paper focuses on aspects of the dataset relevant to the present investigation.

BSETS is a multiday observational study designed to investigate the interplay of sleep and daily experiences. Participants were recruited via branched diffusive convenience sampling, primarily from the families and acquaintances of Semmelweis University students. As BSETS was designed for maximum ecological validity, inclusion criteria were minimal and consisted only of being at least 18 years of age, fluency in Hungarian, and not being recruited from clinical settings. A total of 267 participants provided data for BSETS for a total 1899 days and nights, with variable-specific missingness: descriptive statistics and more precise sample sizes are reported in the Results. The mean age of participants was 28.93 years (SD=12.77 years, range: 18-76), 118 (45.21%) were male and 143 (54.79%) female. 152 participants (59%) were students. Minor deviations of these figures from previous BSETS papers reflect the discovery of new data and corrections.

BSETS participants spent a period of at least 7 days and nights recording their daily experiences using an evening diary, overnight sleep using a Dreem2 mobile EEG device, and their subjective sleep experiences each morning using a second diary. During this period, participants received no intervention and were encouraged not to change their behavior or lifestyle, provided that faithfully recorded their daily and nightly experiences in the diaries.

The Institutional Review Board (IRB) of Semmelweis University, as well as the Hungarian Medical Council (under 7040-7/2021/ EÜIG "Vonások és napi események hatása az alvási EEG-re" [The effect of traits and daily activities and experiences on the sleep EEG]), approved BSETS as compliant with the latest revision of the Declaration of Helsinki. All participants provided written informed consent using a form reviewed and approved by the IRB.

### Food intake

As part of the evening diaries, participants were instructed to recall their food intake from the previous day. Diary entries were produced for predefined meal timings (morning, midday, afternoon, evening and after dinner) and included the time of food consumption (HH/MM), food and beverage item names as described by the participants, subjectively estimated portion sizes, and a subjective satiation rating ranging from 1 to 5. The primary data were collected in handwritten form and manually transcribed into a digital spreadsheet (MS Excel), providing a binary variable indicating the presence of each meal, quantitative variables describing meal timing and satiation and a text variable containing descriptions of the meal and portion size for each meal. Using the primary quantitative meal-level variables, we calculated the following day-level variables:

1. Pre-sleep fasting: the amount of time (in hours) between the last meal and sleep onset
2. Number of meals: the number of meals reported by the participant on a given day
3. Satiation distribution: the sum of the products of each meal’s reported satiation and meal timing, divided by total daily satiation, Σ(satiation*meal time)/Σ satiation. A higher value of this variable indicates that meals providing higher satiation levels were consumed later in the day.

In addition to calculating variables directly from quantitative diary input, we also estimated the energy content (kcal) of each meal based on participants’ textual meal descriptions. For this, we used the large nutritional database of a popular freely available nutrition tracking application (MyFitnessPal, www.myfitnesspal.com). Entries from each food diary were matched to a food item, meal, or recipe in the MyFitnessPal database by a qualified dietitian (LK), the energy value provided by the application was recorded as the estimated energy content of that entry. For each entry, we used the portion sizes reported by participants, or, when unavailable, the default ‘normal’ portion provided by the database. Because this procedure was not standardized, we performed a preliminary validation study (**Supplementary text**), which indicated that energy content could be estimated with reasonable accuracy (r=0.68) compared with a gold-standard method.

### Sleep

Participants wore a Dreem2 mobile EEG headband [32,33] each night. Dreem2 uses dry silicone electrodes to record brain activity at 250 Hz sampling resolution, with a band-pass filter of 0.4-18 Hz. EEG signals were scored using a validated proprietary algorithm [33] to generate the hypnogram, which was then used to obtain the following sleep macrostructure metrics: 1) total sleep time, 2) sleep onset latency, 3) sleep onset time, 4) wake after sleep onset (WASO), 5) sleep efficiency, and 6-9) the percentage of N1, N2, N3 and REM sleep. In line with recent BSETS studies [30,34,35], we log-transformed WASO and sleep onset latency and reflected and log-transformed sleep efficiency. In the case of transformed variables, for the sake of interpretability, we calculate what linear change the effect would imply at the mean level of the dependent variable.

To reduce the number of models, we deviated from previous BSETS studies by omitting less important or redundant macrostructure variables for REM latency, N1-3 and REM duration, number of awakenings, and delta and sigma power from our analyses.

In addition to objective sleep metrics, subjective sleep quality was assessed each morning using the Hungarian version of the Groningen Sleep Quality Scale [36]. The list of nine sleep macrostructure metrics together with subjective sleep quality served as predictors or outcomes, as appropriate, in separate models investigating bidirectional links between food intake and sleep.

### Statistical analysis

The bidirectional associations between sleep and food intake were investigated using multilevel models. Models contained separate within-person and between-person effects, random intercepts per participant, and controls for age, sex, day of the week (weekend/weekday), and lagged outcomes. In models in which the dependent variable was a sleep parameter, we also controlled for time spent awake to distinguish potential effects of food intake from homeostatic effects resulting from extended wakefulness [30,37]. The full model was:

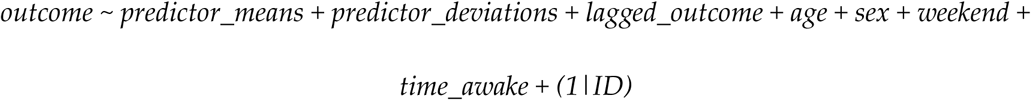

where:

1. “outcome” is either a sleep parameter (in models about food intake predicting subsequent nightly sleep) or a food intake parameter (in models about sleep predicting food intake the subsequent day),
2. “predictor_means” for each participant is the average value of either a sleep parameter or a food intake parameter, estimating between-participant effects,
3. “predictor_deviations” are participant mean-centered values of the predictor (deviations from the individual mean), estimating within-participant effects,
4. “lagged_outcome” is the previous day’s or night’s outcome value, controlling for autoregressive effects,
5. “age”, “sex” and “weekend” are additional covariates,
6. “time_awake” is the number of hours spent awake, an optional covariate in models with a sleep outcome, operationalizing homeostatic effects,
7. “(1|ID)” is a random intercept per participant.

The most important estimand is “predictor_deviations”, that is, the estimated within-person change in sleep following changes in food intake (or vice versa). Although these estimates do not establish causality because time-varying confounding remains possible [38], within-participant estimates have properties which help causal inference. Specifically, by using each participant as their own control they reduce time-invariant confounding by e.g., demographic or psychological variables, due to the time lag helps address reverse causality, and they can falsify causal hypotheses [37].

Between-participant effects (“predictor_means”, correlations of habitual sleep and food intake characteristics) are intended and reported as secondary.

A separate model was fitted for each predictor-outcome combination. The sleep parameters were 1) total sleep time, 2) sleep onset latency, 3) sleep onset time, 4) wake after sleep onset (WASO), 5) sleep efficiency, 6-9): the percentage of N1, N2, N3, and REM sleep, 10) GSQS score, regardless of whether they were predictors or outcomes. Food intake parameters included in all analyses were: 1) number of meals, 2) satiation distribution, 3) total daily energy intake. In models with food intake as the predictor, we additionally included the timing and satiation of the last meal, its energy content, as well as the duration of pre-sleep fasting (4 additional variables). In models using food intake as the outcome, we used the timing and satiation of the first meal, together with its energy content (3 additional variables). First and last meals were defined as whichever meal was reported first/last.

All models with sleep outcomes were corrected for homeostatic effects, except for sleep onset time which is inherently dependent on time spent awake.

Statistical significance was determined as in previous BSETS studies. False discovery rate correction was performed using the Benjamini-Hochberg method, with the corrected significance rate set at p=0.05. The p-values for all predictors of a given outcome were treated as a single family for FDR correction. Additionally, a threshold of tentative significance was set at p=0.01. Effects meeting this threshold but not surviving formal corrections were discussed only when they aided the interpretation of results surviving FDR correction.

All data and code necessary to reproduce the results are available at 10.5281/zenodo.22172516.

## Results

### Descriptive statistics

Full descriptive statistics are reported in **Table 1**.

**Table 1.** Descriptive statistics for all variables in the dataset. For sex, the mean indicates the proportion of male participants. ICC: intraclass correlation coefficient. All variables are reported here in their original units of measurement. Sleep onset time is expressed as hours relative to midnight. The timing and satiation distribution is expressed as time within the day, with 0 as the previous midnight, 0.5 as noon, and 1 as midnight. For convenience, throughout the results, we rescaled regression coefficients to hourly increments of the independent variable.

|  | Valid N | Mean | Median | SD | ICC | Skewness | Kurtosis |
| --- | --- | --- | --- | --- | --- | --- | --- |
| Sex (proportion male) | 261 | 0.45 |  |  |  |  |  |
| Age (years) | 261 | 28.93 | 22.00 | 12.77 |  | 1,27 | 3,36 |
| Total sleep time (min) | 1718 | 396,84 | 402,75 | 91,02 | 0,24 | -0,54 | 3,91 |
| Sleep onset latency (min) | 1718 | 14,91 | 10,50 | 14,85 | 0,31 | 3,62 | 23,96 |
| Sleep onset time (hours relative to midnight) | 1718 | 0,44 | 0,18 | 1,79 | 0,49 | 1,45 | 8,19 |
| Wake after sleep onset (min) | 1718 | 21,02 | 15,00 | 21,56 | 0,38 | 3,53 | 21,35 |
| Sleep efficiency (%) | 1718 | 91,04 | 92,79 | 6,71 | 0,36 | -3,91 | 31,92 |
| N1 percentage (%) | 1718 | 6,08 | 5,59 | 2,49 | 0,52 | 1,41 | 6,24 |
| N2 percentage (%) | 1718 | 46,31 | 46,40 | 9,17 | 0,46 | -0,20 | 3,67 |
| N3 percentage (%) | 1718 | 21,79 | 21,07 | 9,49 | 0,48 | 0,62 | 4,09 |
| REM percentage (%) | 1718 | 25,82 | 25,44 | 7,49 | 0,30 | 0,81 | 9,70 |
| Subjective sleep quality (Likert points) | 1751 | 4,13 | 4,00 | 3,41 | 0,20 | 0,68 | 2,66 |
| Number of meals | 1760 | 3,60 | 4,00 | 0,88 | 0,39 | -0,27 | 2,78 |
| Satiation distribution (time within day) | 1652 | 0,59 | 0,61 | 0,12 | 0,24 | -1,57 | 6,79 |
| Total energy (kcal) | 1757 | 1699,46 | 1608,00 | 667,89 | 0,36 | 0,70 | 3,59 |
| Last meal timing (time within day) | 1525 | 0,85 | 0,83 | 0,09 | 0,35 | 0,32 | 5,43 |
| Last meal satiation (Likert points) | 1644 | 3,91 | 4,00 | 1,18 | 0,28 | -0,85 | 2,76 |
| Last meal energy (kcal) | 1735 | 447,15 | 391,00 | 304,20 | 0,12 | 1,25 | 4,96 |
| Pre-sleep fasting (h) | 1427 | 4,06 | 3,92 | 2,12 | 0,22 | 1,82 | 13,31 |
| First meal timing (time within day) | 1692 | 0,40 | 0,38 | 0,10 | 0,44 | 1,12 | 4,89 |
| First meal satiation (Likert points) | 1704 | 3,85 | 4,00 | 1,08 | 0,37 | -0,79 | 3,05 |
| First meal energy (kcal) | 1739 | 449,55 | 395,00 | 262,03 | 0,25 | 1,13 | 4,77 |
| Wakefulness duration (h) | 1414 | 17,07 | 16,89 | 1,96 | 0,11 | 0,97 | 6,85 |

### Within-person effects of food intake on sleep

Within-person effects represent the primary results of our study. These show an estimated association between deviations in food intake and subsequent deviations in sleep within the same individual. After correcting for multiple comparisons, we identified three significant associations of daytime food intake with subsequent sleep.

First, the timing of the last meal was associated with later sleep onset (within-participant B=0.188 hours per timing hour, p=10^-14^).

Second, longer pre-sleep fasting was associated with later sleep onset (B=0.333 hours per fasting hour, p=10^-54^), reduced total sleep time (B=-4.028 minutes, p=0.003), and reduced sleep efficiency (B=0.01 reflected log units, p=0.006, 0.21% decrease from mean expected per hour). While we report it, the effect on sleep onset is unsurprising because sleep onset is used to calculate pre-sleep fasting and the two variables are highly correlated.

Third, sleep efficiency was improved after higher total daily energy intake (B=-10^-5^ inverted log units per kcal, p=0.005, ∼0.001% increase from mean expected per calorie).

The effects of food intake on subsequent sleep are illustrated on **Figure 1**. Correlations depicted on figures illustrate the strength of associations on a standardized scale.

**Figure 1.**
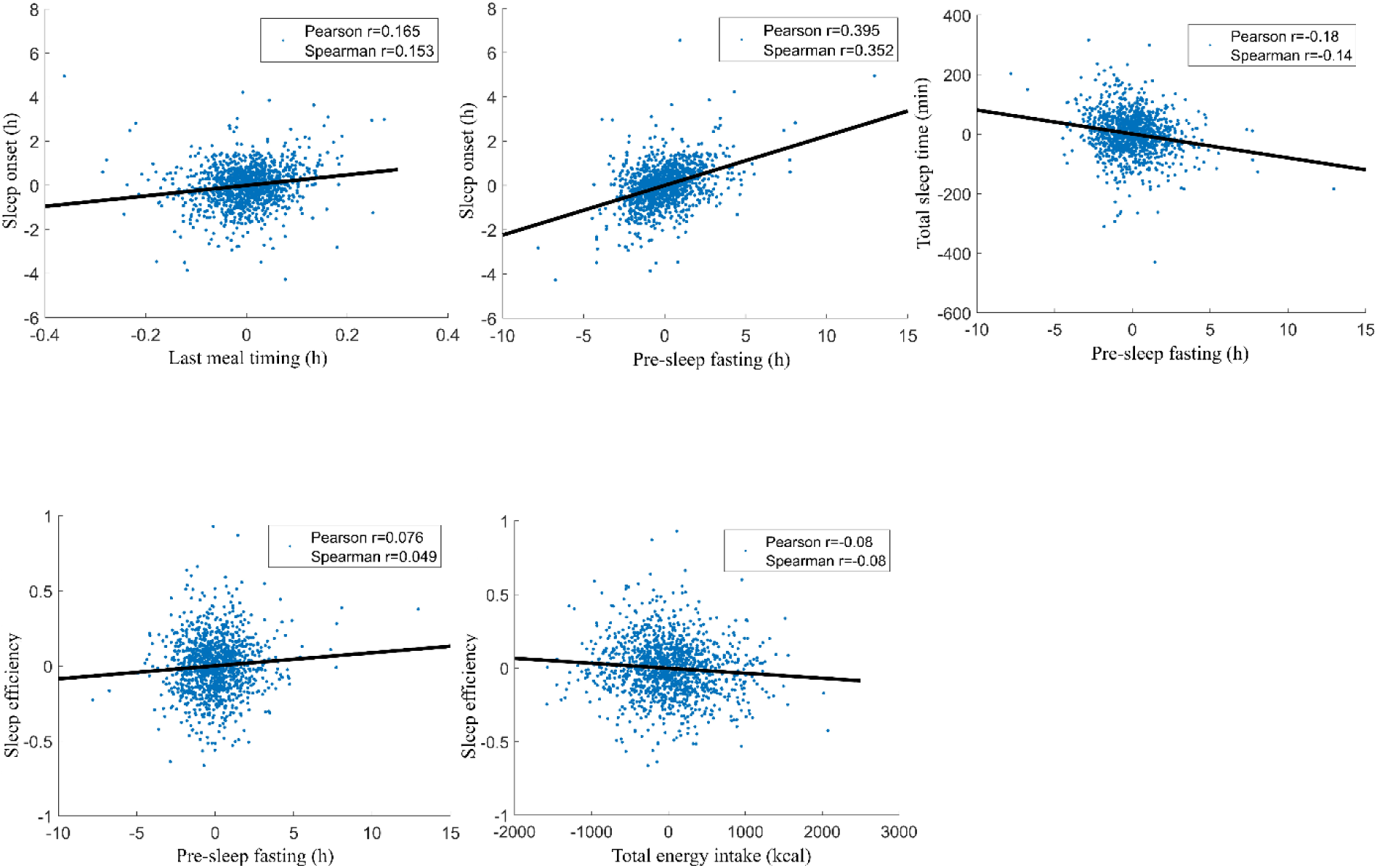
Significant within-participant associations between food intake variables (horizontal axis) and sleep metrics recorded during the subsequent night (vertical axis). The scatterplots show deviations from the individual means, pooled across participants. To illustrate partial correlations net of confounders, control variables (age, sex, day of week, lagged outcomes) were regressed out of deviations. Because the plots show raw residuals, data points are centered around 0. Variable pairs are shown if they reached FDR-corrected significance. Both Pearson and Spearman partial correlations are shown to illustrate the effect of outliers on the associations.

### Effects of sleep on food intake

Three sleep metrics, total sleep time, sleep onset time, and subjective sleep quality had significant associations with next-day food intake.

Increased total sleep time was associated with a greater number of meals (B=0.48 meals per hour, p=0.006) and increased satiation (B=0.001 Likert points, p=0.002) with the first meal.

Later sleep onset was, conversely, associated with fewer next-day meals (B=-0.069 meals per hour, p=10^-5^), as well as later timing (B=0.336 hours, p=10^-14^) and increased energy (B=21.12 kcal, p=10^-4^) intake during the first meal. Furthermore, the temporal distribution of self-reported satiation shifted to a later time (B=0.216 hours, p=0.001).

Lower self-rated sleep quality was associated with earlier timing of (-0.048 hours per GSQS point, p=0.002) and reduced satiation with (B=-0.045 Likert points, p=0.009) the first meal.

The effects of sleep on next-day food intake are illustrated in **Figure 1**.

**Figure 1.**
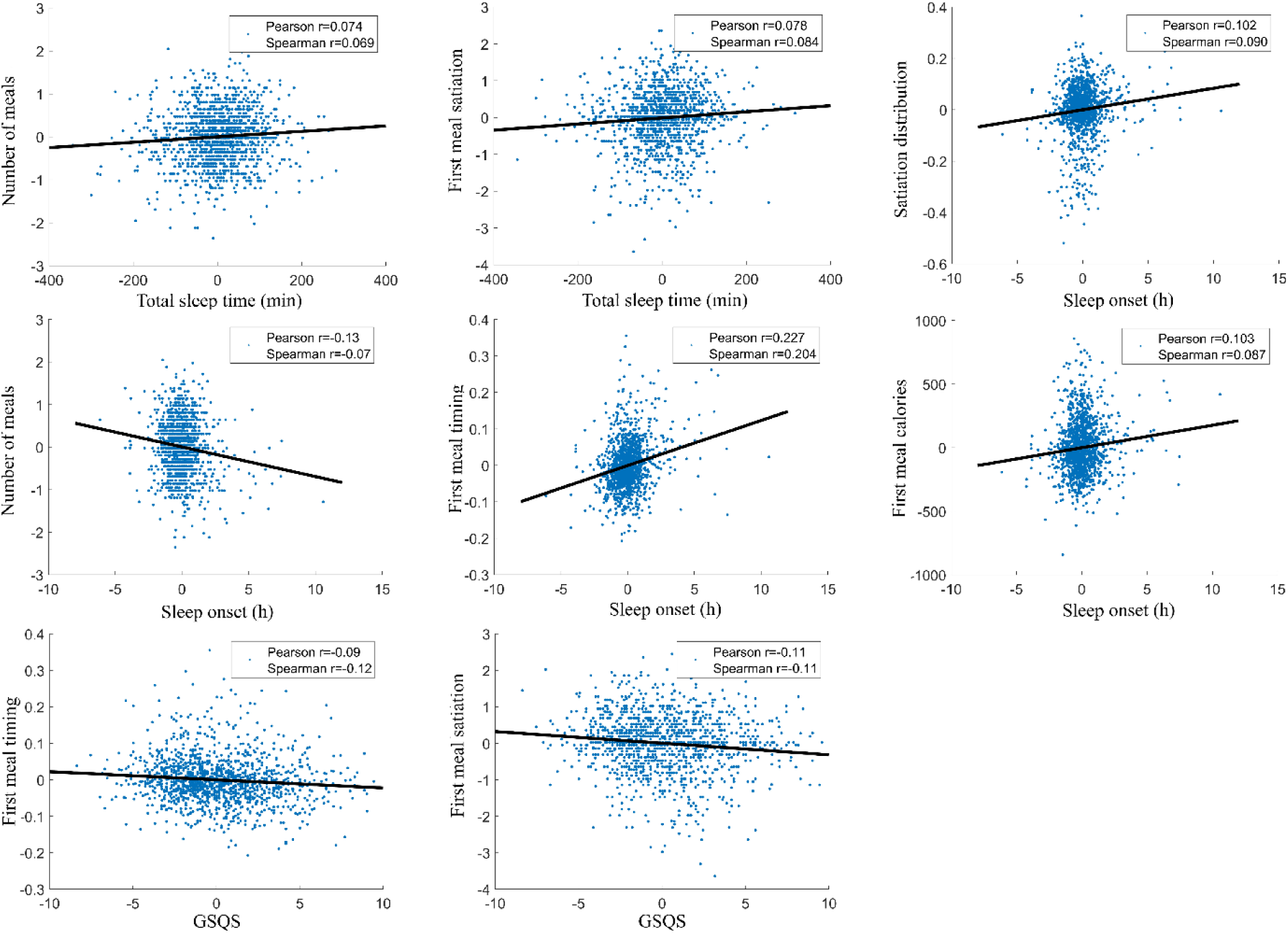
Significant within-participant associations between sleep metrics (horizontal axis) and food intake variables recorded during the next day (vertical axis). The scatterplots show deviations from the individual means, pooled across participants. To illustrate partial correlations net of confounders, control variables (age, sex, day of week, lagged outcomes) were regressed out of deviations. Because the plots show raw residuals, data points are centered around 0. Variable pairs are shown if they reached FDR-corrected significance. Both Pearson and Spearman partial correlations are shown to illustrate the effect of outliers on the associations. GSQS: subjective sleep quality recorded by the Groningen Sleep Quality Scale.

**Figure 2.**
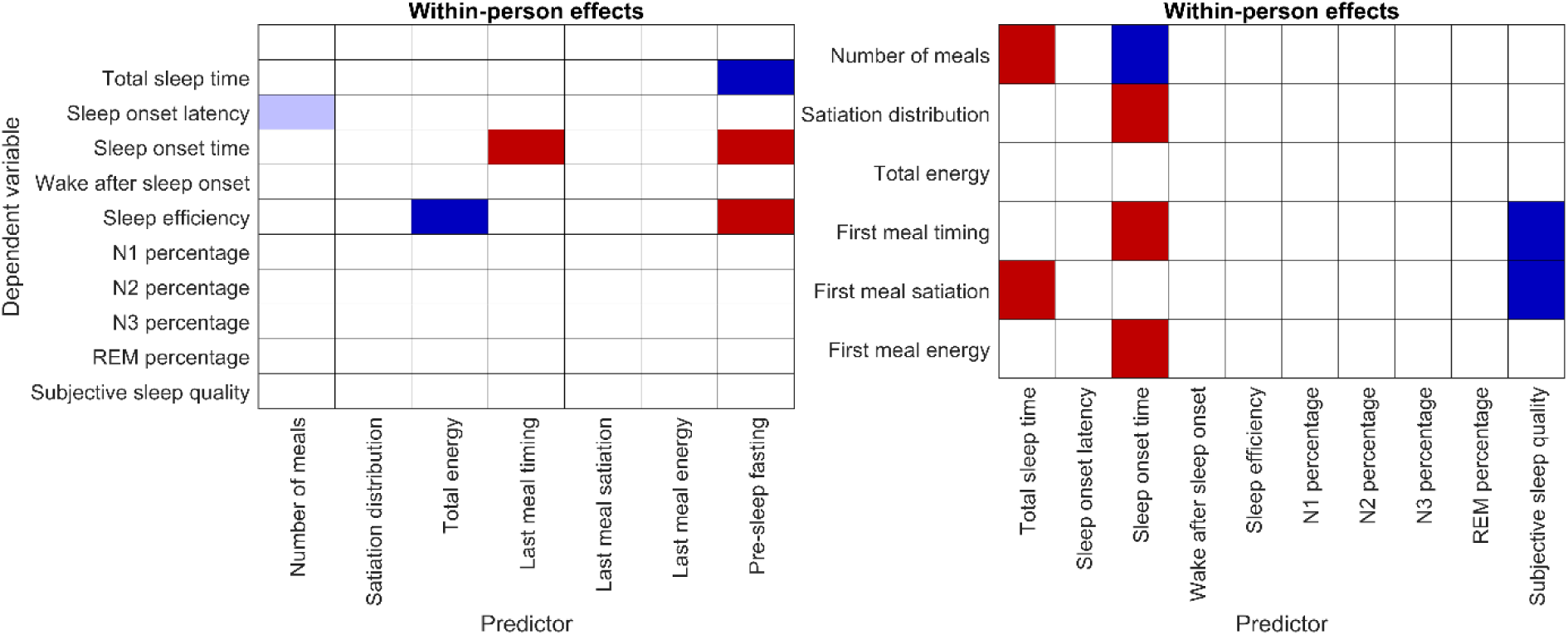
summarizes all bidirectional within-person effects. A visual summary of all within-person effects in the food intake-to-sleep (left) and sleep-to-food intake (right) direction. The matrices show all predictor-outcome combinations, with predictors on the horizontal and outcomes on the vertical axis. Cells are shaded with a light color if tentative significance (p<0.01) was passed, and dark color if the effect survived full FDR correction. Blue cells indicate negative and red cells positive associations. Note that due to the transformation of this variable, negative effects on sleep efficiency indicate an improvement and positive effects a decline. All models are corrected for age, sex, day of the week, lagged outcomes, and, in the case of food intake-to-sleep models, homeostatic effects.

### Between-participant effects

Between-participant effects also showed a link between the habitual timing of sleep and food intake. Both the timing of the first (B=5.92 minutes per hour, p=10^-18^) and the last (B=10.52 minutes per hour, p=10^-21^) meal was positively associated with sleep onset time, as was the daily distribution of satiation (B=0.48 hours per hour, p=10^-11^). Longer habitual pre-sleep fasting was associated with shorter total sleep time (B=-7.069 minutes per hour, p=^10-4^) and later sleep onset (B=0.289 hours per hour, p=10^-6^). These findings are illustrated on **Supplementary figure S1**.

Full outputs are available at 10.5281/zenodo.22172516

## Discussion

In this study, we investigated bidirectional associations between food intake and sleep in an ecologically valid, within-participant design. Our findings suggest that, in line with previous literature [15], bidirectional associations exist even in the absence of interventions. With some exceptions, associations primarily involved the timing of sleep and food intake, rather than other measured characteristics of sleep or energy intake, suggesting that both primarily serve as time cues for the other.

We found that food intake primarily affects the timing, and not the structure or quality, of subsequent sleep. Later meals were followed by later sleep onset times, an effect apparent both in within- and between-participant analyses. These results support the important role of food intake as a temporal cue [13,39], and are in line with findings from naturalistic [15] and interventional [40] studies. The timing of meals may serve as a pacesetter for the circadian clock, with not only habitually delayed eating associated with delayed sleep, but the same person’s sleep expected later after a later cessation of food intake. This pattern of findings also coheres with previous BSETS literature [30] which found that daily experiences other than food intake also primarily affect the timing, and not the structure or quality of sleep.

Some findings also supported a somnogenic role of food intake. Specifically, we found improved sleep efficiency after greater daily energy intake, and reduced sleep efficiency if pre-sleep fasting was longer. Refraining from eating too much or too late often appears frequently in sleep hygiene recommendations even by reputable sources ^1^, although evidence for this claim is mixed, with only a few small interventional studies conducted. At least one study indeed found a negative effect of late eating on sleep quality and other sleep macrostructure variables [41], but some found no effect [39,42], and at least one positive effect [43]. No effect was found in the single naturalistic study other than our own [15]. In the light of this heterogeneous literature, our finding of a positive effect of late eating should not be considered definitive, the topic warrants further research. We note, however, that sleep hygiene recommendations against late eating are based on little evidence and warrant further evaluation.

In the other temporal direction, we also found an effect of the timing and duration of sleep on the timing and energy content of next-day food intake. After nights with later sleep onset, first meals were consumed later and contained a greater amount of energy, presumably because they were more often intended as midday meals rather than breakfast. The temporal distribution of satiation also shifted later, and the number of meals was reduced with no effect on energy intake, consistent with habitual food intake compressed into a shorter, later feeding window. Longer sleep was associated with more meals and greater satiation with the first meal, with no effect on total energy intake.

A large literature of interventional studies supports the negative effect of sleep restriction on the quality of next-day food intake [23,26,44,45]. Such studies are scarce due to the difficulty of manipulating sleep timing, with only a single such study of 5 participants [46] identified in a recent meta-analysis [23]. However, many cross-sectional studies support an association between early chronotype and healthy eating patterns [47–49], although these correlational studies are subject to confounding. In our naturalistic study, we did not find evidence for changes in energy intake after either longer or later sleep: in line with results from the single other available observational study [15], we found that both variables are primarily associated with a temporal redistribution of next day food intake and satiation, mirroring the effects of food intake on sleep in the other temporal direction. This discrepancy is most likely explained by the fact that almost all (e.g., in a recent meta-analysis [23], all but four) interventional studies restrict sleep for several days, studying the effects of chronic sleep restriction, whereas naturalistic studies effectively investigate the effects of a single night. In combination, these results suggest that while energy intake is increased after chronic sleep restriction, making the latter a risk factor for obesity, it is resilient to acute effects. However, further possible explanations for the lack of an effect on energy intake may be the limited precision of energy measurements in observational studies, and the lack of data on diet quality, and possible nonlinear effects of the amount of sleep restriction on food intake, which are apparent following the pronounced interventions of experimental studies but are not triggered by the relatively modest normal day-to-day variation in sleep duration. Further research is needed to determine if the timing of sleep is a modifiable behavioral interventional supporting healthier next-day eating patterns, and whether low levels or fewer days of sleep restriction fails to trigger changes in food intake.

Interestingly, subjective sleep quality was associated with the timing and satiation of the first meal, even though they were reported a full wakeful period apart in two separate diaries. In other words, if participants reported better subjective sleep quality in the morning, they ate later and recalled their first meal as more satiating several hours later, in the evening diary. One possible explanation is that positive perceptions of sleep influence the perception of meals consumed shortly after. An interesting alternative possibility is that a stable baseline level of mood (“affective inertia” [50]) is already present immediately after waking up; this, in turn, causes a correlated bias in all subjective reports created during that day, including subjective sleep quality in the morning and retrospectively reported meal satiation in the evening. Our research design does not conclusively establish the correct explanation for this pattern, which therefore needs replication and further investigation.

Our work has limitations. Most importantly, we used a novel technique to record energy intake which, although performing favorably in our validation study, is not perfectly exact. Errors in the evaluation of food intake may have led to false negative findings. In the future, AI-vision based evaluation of meal images may provide more accurate estimates [51]. Second, we used a relatively limited set of sleep parameters derived from mobile EEG recordings. It is possible that more subtle characteristics of sleep affect (or are affected by) food intake. Finally, a general limitation of all BSETS studies is the oversampling of young, healthy participants from relatively high socioeconomic status, which limits the generalizability of findings to clinical or elderly samples where sleep complaints are the most prevalent.

In conclusion, we found modest but significant bidirectional relationships between food intake and sleep. In both directions, food intake and sleep primarily served as a temporal cue for the other. Contrary to published recommendations, we found no evidence that reducing late-night eating is beneficial for sleep.

## Data availability

Data and code to reproduce analyses are available at 10.5281/zenodo.22172516.

## Footnotes

1 https://www.mayoclinic.org/healthy-lifestyle/adult-health/in-depth/sleep/art-20048379 https://www.health.harvard.edu/healthy-aging-and-longevity/sleep-hygiene-simple-practices-for-better-rest https://www.thensf.org/get-healthy-sleep-by-eating-right-on-schedule https://www.ama-assn.org/public-health/prevention-wellness/8-things-doctors-want-patients-know-about-healthy-sleep-habits

