## Supplementary Material for "Bidirectional links between food intake and sleep: a naturalistic mobile EEG study"

**Supplementary text**

We performed a preliminary validation study to ensure that the nutritional values returned by the method described in the main paper (henceforth „BSETS method") were valid estimates of the true nutritional value of meals consumed by participants.

In the validation study, we recruited three registered dietitians and three additional participants to participate in a one-week procedure. Participants downloaded the MyFitnessPal application, and they were requested to weigh and record each meal they had using precise gram weights. These records were considered to be the gold standard. Additionally, they were asked to recall their food intake in evening diaries identical to those used in BSETS. They were familiarized with the content of a sample of actual BSETS diaries and asked to report food intake in similar detail instead of trying to recall the precise nutritional values they recorded during the day.

The content of the evening diaries was processed using the standard BSETS method reported in the main paper. Thus, we obtained quantitative estimates of nutrients using two independent methods: using precise scale-assisted entries by the participants themselves as the gold standard, and the processing of evening food intake diaries using the BSETS method.

The six participants recorded 149 meals in total. While MyFitnessPal returns data on a large number of macro- and micronutrients, the availability and quality of data on these in the database on some was variable. Therefore, we focused our analysis on energy intake (kcal), a key variable with high availability. Because we were interested in the association of energy intake with other variables, which is unaffected by biases in the mean, we operationalized validity the Pearson correlation with the gold standard.

Based on this analysis, the validity of energy estimates recorded by the BSETS methods was r=0.68. Raw data generated using the validation study is available at 10.5281/zenodo.22177284.

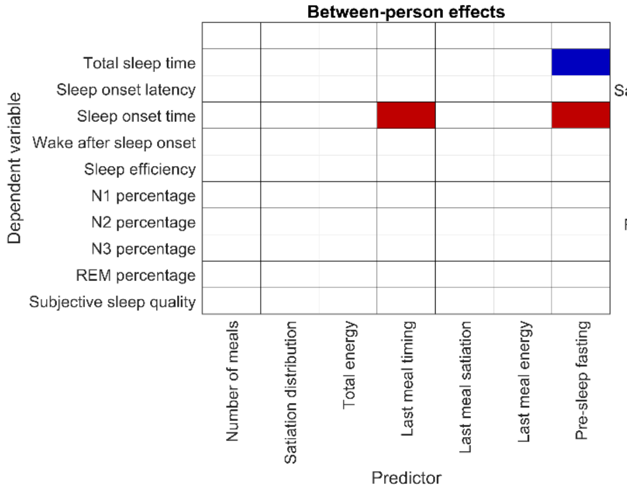


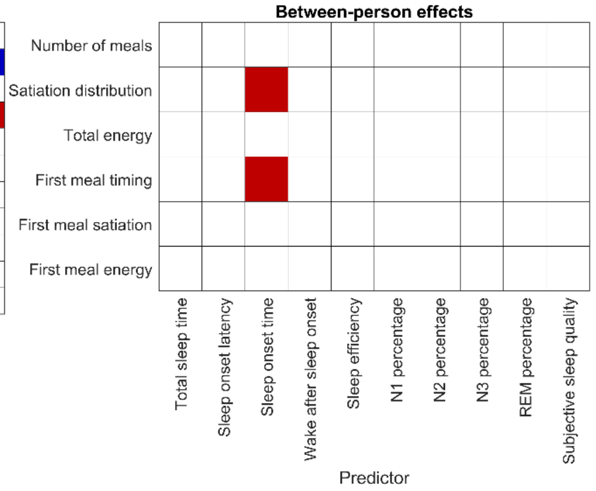

**Supplementary figure S1.** A visual summary of all between-person effects from food intake-to-sleep (left) and sleep-to-food intake (right) models. The matrices show all predictor-outcome combinations, with predictors on the horizontal and outcomes on the vertical axis. Cells are shaded with a dark color if the effect survived full FDR correction. Blue cells indicate negative and red cells positive associations. All models are corrected for age and sex.
